# Magnitude of road traffic accident related injuries and fatalities in Ethiopia

**DOI:** 10.1101/382333

**Authors:** Teferi Abagaz, Samson Gebremedhin

## Abstract

**Background:** In many developing countries there is paucity of evidence regarding the epidemiology of road traffic accidents (RTAs). The study determines the rates of injuries and fatalities associated with RTAs in Ethiopia based on the data of a recent national survey.

**Methods:** The study is based on the secondary data of the Ethiopian Demographic and Health Survey conducted in 2016. The survey collected information about occurrence injuries and accidents including RTAs in the past 12 months among 75,271 members of 16,650 households. Households were selected from nine regions and two city administrations of Ethiopia using stratified cluster sampling procedure.

**Results:** Of the 75,271 household members enumerated, 123 encountered RTAs in the reference period and rate of RTA-related injury was 163 (95% confidence interval (CI): 136-195) per 100,000 population. Of the 123 causalities, 28 were fatal, making the fatality rate 37 (95% CI: 25-54) per 100,000 population. The RTA-related injuries and fatalities per 100,000 motor vehicles were estimated as 21,681 (95% CI: 18,090-25,938) and 4,922 (95% CI: 3325-7183), respectively. Next to accidental falls, RTAs were the second most common form of accidents and injuries accounting for 22.8% of all such incidents. RTAs contributed to 43.8% of all fatalities secondary to accidents and injuries. Among RTA causalities, 21.9% were drivers, 35.0% were passenger vehicle occupants and 36.0% were vulnerable road users including: motorcyclists (21.0%), pedestrians (12.1%) and cyclists (2.9%). Approximately half (47.1%) of the causalities were between 15-29 years of age and 15.3% were either minors younger than 15 years or seniors older than 64 years of age. Nearly two-thirds (65.0%) of the victims were males.

**Conclusion:** RTA-related causalities are extremely high in Ethiopia. Male young adults and vulnerable road users are at increased risk of RTAs. There is a urgent need for bringing road safety to the country’s public health agenda.

## Background

Road traffic accidents (RTAs) are huge public health and development problems. Every year nearly 1.3 million people lose their lives on the road and as many as 50 million others are injured [1]. Globally 17 road fatalities per 100,000 population per annum are reported. RTA is the second leading cause of death in economically active population group of 15-44 years of age and more than 75% of RTA casualties occur in this age group [2–4]. In many countries the estimate economic loss due to RTAs is as high as 3% of their gross domestic products [5].

The burden of RTA is disproportionally high in low- and middle-income countries (LMIC) were over 85% fatalities and 90% of disability-adjusted life years lost are reported [3,6,7]. Fatalities secondary to RTAs is more than double in LMIC than in high-income countries. Globally RTA fatalities remain more or less constant since 2007; yet, in many developing countries the rates are increasing [8]. Especially Africa faces the highest annual rate of road fatalities in the world – 27 per 100,000 population [8]. In the next few decades, the problem can even soar due to the ongoing rapid economic growth and increase in motorization in the continent [2,6,9].

Despite the growing burden of RTAs, road safety remains a neglected issue in many developing countries and the health sector has been slow to recognize it as a priority public health problem [8,10]. A large body of evidence suggests that RTAs are easily preventable and many high income countries have successfully reduced the incidence through proven and cost-effective interventions [2,11]. The Sustainable Development Goals (SDGs) Goal-3 sets an ambitious target to halve the global number of fatalities and injuries from RTAs by 2020 [12].

Ethiopia as many African countries is facing enormous road safety crisis. Each year thousands of road users are killed and the majority of them are economically active population [13].

According to the estimate of the World Health Organization (WHO), the prevalence of road traffic fatality in Ethiopia for the year 2013 was 25.3 per 100,000 population and the rate is among the highest in the world [8]. Factors contributing to the high incidence of RTAs in Ethiopia include rampant reckless driving behaviors, poor road network, substandard road conditions, failure to enforce traffic laws and poor conditions of vehicles [14].

In many developing countries including Ethiopia there is paucity of evidence regarding the incidence of RTA-related injuries and fatalities. Further, the available estimates based on official reports are likely to underestimate the extent of the problem [15]. In this study, we examined the magnitude of RTA-related injuries and fatalities in Ethiopia based on the secondary data of nationally representative survey conducted in 2016.

## Methods

### Study design

This study is based on the data of the Ethiopian Demographic and Health Survey (EDHS) conducted in 2016. The DHS is a standard survey that has been carried out in several LMIC including Ethiopia by national agencies in close collaboration with the DHS program. The DHS gathers nationally representative health and population data and provides updated estimates of key health and demographic indices. The EDHS – 2016 for the first time has collected information on injuries and accidents including RTAs [16].

### Sampling approach

The EDHS 2016, collected information about the occurrence of road traffic and other accidents in the preceding 1 year of the survey among 75,271 members of 16,650 households. As the study was conducted based on readily available data, we did not make priory sample size calculation.

The EDHS – 2016 employed stratified two-stage cluster sampling procedure designed to provide representative sample for multiple health and population indicators at national, place of residence (urban-rural) and sub-national levels (nine geographical regions and two city administrations). Initially, 645 enumeration areas (EAs) (202 in urban areas and 443 in rural areas) were drawn using probability proportional to size (PPS) sampling approach from a complete list of 84,915 EAs defined in the recent 2007 population census. Then in every selected EA an exhaustive listing of households was made and 28 households were selected using systematic random sampling approach. In the selected households, enumeration of the entire members was made and information about the occurrence of injuries and accidents including RTAs among all household members was collected primarily from the household head or his/her partner [16].

### Data collection

The EDHS data were collected from January to June 2016 using standardized and pretested questionnaires prepared in three major local languages. The data collectors, editors and supervisors received a four-weeks training [16]. Information about the occurrence of injuries and accidents including RTAs was assessed by asking the respondent whether any child or adult in the household was killed or injured in the past 12 months in any accident/injury severe enough that the victim could not carry out their normal activities for at least a day [16]. When such incidents were reported, additional information about the cause, type, length of injury, whether the injury was fatal or not, characteristics of the victim (age, sex) were explored.

### Data analysis

The “individual” and “household” datasets of the DHS 2016 survey were downloaded separately from the DHS Program website [17] in SPSS format. Information about the victims were extracted from the “household” dataset, appended to the “individual” dataset and duplicate observations were removed. In order to accommodate for the complex sampling design employed in the survey, weighted data analysis was employed. Data weights were computed using sampling weights provided in the dataset and post-stratification weight developed based on the 2016 population size of the nine regions and two city administration of the country [18]. The analyzed dataset is available from: https://dhsprogram.com/Data/.

The rates of injuries and fatalities secondary to RTAs are presented per 100,000 population. The rates were also converted and provided per 100,000 vehicles; inconsideration of the 2016 midyear population size of the country and the registered number of motor vehicles in country in the same year. Ninety-five percent confidence intervals (95% CI) were estimated for the rates using the binomial CI calculator of the STATA software. Household wealth index, a composite indicator of household wealth status, was determined using Principal component Analysis (PCA) based on the ownership of selected valuable household assets. Association between RTA and basic socio-demographic characteristics (age, sex, place of residence, household wealth index) was determined using Bivariable Binary Logistic Regression analyses and the outputs are provided using Odds Ratio (OR) with the respective 95% CI. Multivariable logistic regression analysis was not employed because we did not expect possible confounding among independent variables considered in the study.

### Ethical considerations

The EDHS 2016 study was ethically cleared by the National Research Ethics Review Committee, Ministry of Science and Technology of Ethiopia [16]. Data were collected after taking informed consent from the respondents.

## Results

### Socio-demographic characteristics

The EDHS 2016 survey gathered data from 16,650 households across the nine regions and two city administration of the country. The weighted sample distribution shows 80.4% of the households were drawn from rural areas while the remaining 19.6% were from urban areas. The mean (±SD) age of the respondents was 40.1 (±16.1) years and the majorities (57.1%) were women. More than half of the respondents (56.7%) had no formal education and very few (6.1%) had attained tertiary level of education. The median (inter-quartile range) of the household size was 4 (3-6) (Table 1).

**Table 1:** Socio-demographic characteristics of the respondents, Ethiopian Demographic and Health Survey, 2016.

| Respondent's characteristics (n=16,650) | Frequency | Percentage |
| --- | --- | --- |
| Age in years | 2,553 | 15.3 |
| 15-24 | 4,817 | 28.9 |
| 25-34 | 3,409 | 20.5 |
| 35-44 | 2,412 | 14.5 |
| 45-54 | 1,868 | 11.2 |
| 55-64 | 1,590 | 9.5 |
| 64+ | 2,553 | 15.3 |
| Sex |  |  |
| Male | 7,037 | 42.3 |
| Female | 9,613 | 57.7 |
| Educational status |  |  |
| No formal education | 9,439 | 56.7 |
| Primary | 4,844 | 29.1 |
| Secondary | 1,357 | 8.1 |
| Higher | 1,010 | 6.1 |
| Place of residence |  |  |
| Urban | 3,259 | 19.6 |
| Rural | 13,391 | 80.4 |
| Number of household members |  |  |
| Less than 4 | 5,675 | 34.1 |
| 4-5 | 5,294 | 31.8 |
| 5 or above | 5,682 | 34.1 |

### Magnitude of road traffic accident related injuries and fatalities

The occurrence of RTAs in the past 12 months of the survey was assessed among 75,271 members of the 16,650 households included in the survey. It was found that 123 road accidents – equivalent to a rate of 163 (95% CI: 136-195) per 100,000 population – occurred in the reference 12 months period. Of the 123 causalities, 28 were fatal injuries, making the road traffic fatality rate 37 (95% CI: 25-54) per 100,000 population.

In addition to RTAs, the survey collected information about the occurrence of other forms of accidents and injuries in the same reference period including violence/assault, burn, accidental fall, drowning and poisoning. The rate of all forms of injuries and accidents including RTAs was 716 (95% CI: 657-779) per 100,000 population. RTAs were the second most common form of accidents and injuries contributing to 22.8% of all such incidents. The other frequent forms of accidents and injuries were fall (33.2%), violence/assault (14.1%), burns (9.2%), bite or kick by animals (3.9%), poisoning (2.0%) and drowning (1.5%). Nearly one-in-ten (11.9%) of all the injuries and accidents were fatal. RTAs contributed to 43.8% of all these fatalities.

Among 123 individuals who sustained RTAs in the reference period, 21.9% were drivers and 35.0% were passenger vehicle occupants. Further, 36.0% of the victims were vulnerable road users: motorcyclists (21.0%), pedestrians (12.1%) and cyclists (2.9%). Young adults 15 and 29 years constitute 47.1% of all road accidents; further, 15.3% of the casualties were either children younger than 15 years or seniors older than 64 years of age. Two-thirds (65.0%) of the victims were males.

Among 95 RTA survivors, the duration that the victim was excluded from engaging in normal activities secondary to the injury was assessed. Nearly half (46.2%) of the causalities were precluded from their usual duties for more than a month (Table 2).

**Table 2:** Characteristics of the road traffic accident victims, Ethiopia, 2016.

| Characteristics of the road traffic accident victims (n=123) | Frequency | Percent |
| --- | --- | --- |
| Type of the victim (n=123) |  |  |
| Passenger vehicle occupants | 43 | 35.0 |
| Drivers | 27 | 21.9 |
| Motorcyclists | 26 | 21.0 |
| Pedestrian | 15 | 12.1 |
| Cyclists | 4 | 2.9 |
| Others | 8 | 6.9 |
| Sex (n=123) |  |  |
| Male | 80 | 65.0 |
| Female | 43 | 35.0 |
| Age in years (n=123) |  |  |
| 0-14 | 14 | 11.5 |
| 15-29 | 58 | 47.1 |
| 30-64 | 46 | 37.6 |
| 65 years or above | 5 | 3.8 |
| Place of residence (n=123) |  |  |
| Urban | 37 | 30.4 |
| Rural | 86 | 69.6 |
| Duration that the RTA survivor was precluded from regular activities due to the injury (n=95) |  |  |
| Less than 7 days | 15 | 16.3 |
| Between 8 to 30 days | 35 | 37.3 |
| Between 1 to 6 months | 34 | 36.1 |
| More than 6 months | 10 | 10.1 |

The rate of RTA related injuries and fatalities provided per 100,000 populations can also be per 100,000 vehicles. Accordingly, the RTA related injuries and fatalities per 100,000 motor vehicles were 21,681 (95% CI: 18,090-25,938) and 4,922 (95% CI: 3325-7183), respectively.

### Socio-economic factors associated with RTAs

Comparison of the level of occurrence of RTAs across selected socio-demographic characteristics suggested statistically significant variations by place of residence, age, sex and household wealth index. Urban residents had 2.5 times increased odds of the accident as compared to rural residents. Males were nearly at two times more likely to sustain road accidents than females. As compared to children 0-14 years of age, adults aged 15-29, 30-64 and 65 years or above had eight, six and four times increased odds of RTAs, respectively. Similarly, the occurrence of RTAs was more frequent among individuals from better-off households (Table 3).

**Table 3:** Association between selected socio-demographic factors and road traffic accident, Ethiopia, 2016.

| Variable | RTA |  | Odds ratio (95% CI) |
| --- | --- | --- | --- |
|  | Yes | No |  |
| Place of residence |  |  |  |
| Urban | 37 | 11,351 | 2.45 (1.67-3.60)* |
| Rural | 86 | 63,797 | 1 |
| Sex |  |  |  |
| Male | 80 | 37,132 | 1.97 (1.32-2.76)* |
| Female | 43 | 38,016 | 1 |
| Age (years) |  |  |  |
| 0-14 | 14 | 35,334 | 1 |
| 15-29 | 58 | 17,453 | 8.26 (4.62-14.76)* |
| 30-64 | 46 | 19,003 | 6.06 (3.34-10.98)* |
| 65 years or above | 5 | 3,332 | 3.51 (1.24-9.96)* |
| Wealth index (quintile) |  |  |  |
| Lowest | 14 | 16,561 | 1 |
| Second | 22 | 14,657 | 1.74 (0.90-3.38) |
| Middle | 13 | 14,621 | 1.01 (0.48-2.15) |
| Fourth | 29 | 14,791 | 2.22 (1.18-4.19)* |
| Highest | 45 | 14,518 | 3.56 (1.97-6.45)* |

## Discussion

This study based on nationally representative and population-based data indicated that the rates of injuries and fatalities secondary to RTAs are exceptionally high in Ethiopia and more than one-third of such accidents occur in vulnerable road users including motorcyclists, pedestrians and cyclists. Further, RTAs are more frequent among males, adults 15-29 years of age, urban residents and individuals from better-off households.

In 2013 WHO estimated on yearly basis 25.3 road accident fatalities per 100,000 population occur in Ethiopia [8]. The figure is within the 95% confidence limits of our estimate; 37 (95% CI: 25-54) per 100,000 population. Yet, in addition to sampling variation, the higher point estimate observed in our study can be explained by methodological variations. In this study RTA fatality was assessed by asking the respondent whether any member of the household was killed in the past 12 months due to road accident or not. However WHO counts deaths that exclusively occur within 30 days of the accident [8]. Accordingly, latter is likely to exclude late fatalities. Further, WHO estimates are made based on official national reports and a couple of studies testified that official reports are likely to underestimate the magnitude of the RTAs [6,19]. A population based study in Dar es Salaam, Tanzania found that official reports were only filed for about half of the actual RTAs [19].

While more than two-thirds of all traffic accidents occur among rural residents, urban residents have an elevated injury rate. This is likely due to the higher volume of traffic and human mobility in urban areas. To the best of our knowledge, no study explored the urban – rural variations in the incidence of RTAs in developing countries. However, unlike our finding, two studies conducted in developed countries suggested that both crash injury and fatality rates are higher in rural than in urban areas [20, 21]. Here it is also important to note that we compared the urban – rural differences in the occurrence of RTAs based on the usual place of residence of the victim, not on the actual location of the incident.

We found that economically active population 15-64 years of age contributes to nearly 85% of the RTA burden. Especially adults 15-29 years take the highest toll. This finding is suggestive of the high loss of productivity associated with road accidents in Ethiopia. A study conducted in Kenya also found that more than three quarters of all road traffic victims were economically productive young adults [4]. Similarly, WHO estimated that adults aged 15-44 years account for nearly half of the global road traffic deaths [5]. These segments of population are usually the breadwinners of the family and they might be expected to travel long distances to fulfill the needs of their families.

Most of the studies conducted so far consistently indicated pedestrians are the main victims of RTAs. In LMIC pedestrians may account up to 75% of the total RTA fatalities [3,6]. However, in this study car occupants were more frequently involved in road crashes. This might be explained by the fact that most of the previous studies were based on institutional data and they were mainly limited to urban settings. In urban areas of low income countries pedestrians commonly share road with fast moving vehicles as a result they may disproportionately sustain RTAs. However, in our study 80% of participants were rural residents with relatively less exposure to RTA as a pedestrian. A couple of studies have suggested that passengers are at higher risk of sustaining injury in intercity highways [4,22]. In Libya a study based on medical records indicated vehicle occupants were admitted more often than pedestrians [23].

The study indicated two-thirds of the victims of RTAs are males and they have nearly twofold increased odds of RTAs than females. Previous studies from both in developed and developing countries also concluded that males are disproportionately involved in traffic accidents [24–28]. Studies also documented clear variations in the accident involvement and risky driving behaviors between the two genders in favor of females [25,29–31]. Though males are frequently considered as skilled drivers, they often engage in risky behaviors including trespassing speed limits, reckless overtaking, non-use of seatbelts and driving under the influence of alcohol [25,29]. Male drivers also tend to have lower attention, impatience and risk perception than females [25,30]. The more frequent involvement of males in cycling and riding of two-wheeled motor vehicles may also contribute for their disproportionate burden [24]. Further, in developing countries like Ethiopia, females are less engaged in outdoor activities and this may reduce their susceptibility to RTAs.

The typical strength of the study is the fact that it availed nationally representative information on the magnitude of both RTA-related injuries and fatalities based on population based data.

Most of the existing estimates only measured incidence of fatalities and they were reliant on official reports which are liable to coverage errors [6,19]. The study also assessed the epidemiological significance of RTAs in comparison with other forms of accidents and injuries. Such information has not been frequently presented in previous undertakings.

Conversely, the following key limitations should be considered while interpreting the findings of the study. Firstly, despite the large sample size we used, rates of RTA-related injuries and fatalities were estimated based on small numbers of accidents. This has undoubtedly affected the precision of the estimates. Further, the smaller numbers of events did not allow us for estimation of rates for narrower age intervals and different geographical regions of the country. Secondly, the DHS only enumerates RTAs that were severe enough to preclude the victim out their normal activities for at least a day. While the definition is sensible enough to detect moderate injuries, it may exclude minor accidents like soft tissue injuries. Finally, the study only assessed the sociodemographic differentials of RTAs and did not explore more critical risky behaviors of drivers.

## Conclusion

Road accident related injuries and fatalities are extremely high in Ethiopia. Males, adults 15-29 years of age, individuals from better-off households and vulnerable road users including motorcyclists, pedestrians and cyclists are at increased risk of RTAs. RTA should be considered as a priority public health problem in Ethiopia and there is a need for bringing road safety to the country’s public health agenda. While safety interventions including law enforcements and education should include all road users; implementing interventions targeted at young adults and vulnerable road users is likely to be helpful.

## Acknowledgement

The authors acknowledge Measure DHS for making the Ethiopia DHS 2016 dataset accessible for use.

